## Appendix for "Genome-scale metabolic modeling reveals SARS-CoV-2-induced metabolic changes and antiviral targets"

**Table of Contents**

**Appendix Figures**

Appendix Figure S1. Comparison of the differentially expressed (DE) genes upon SARS-CoV-2 infection across datasets with different DE cutoffs.

Appendix Figure S2. Pathway enrichment analysis of SARS-CoV-2-induced differential changes with the alternative differential expression algorithm limma-voom.

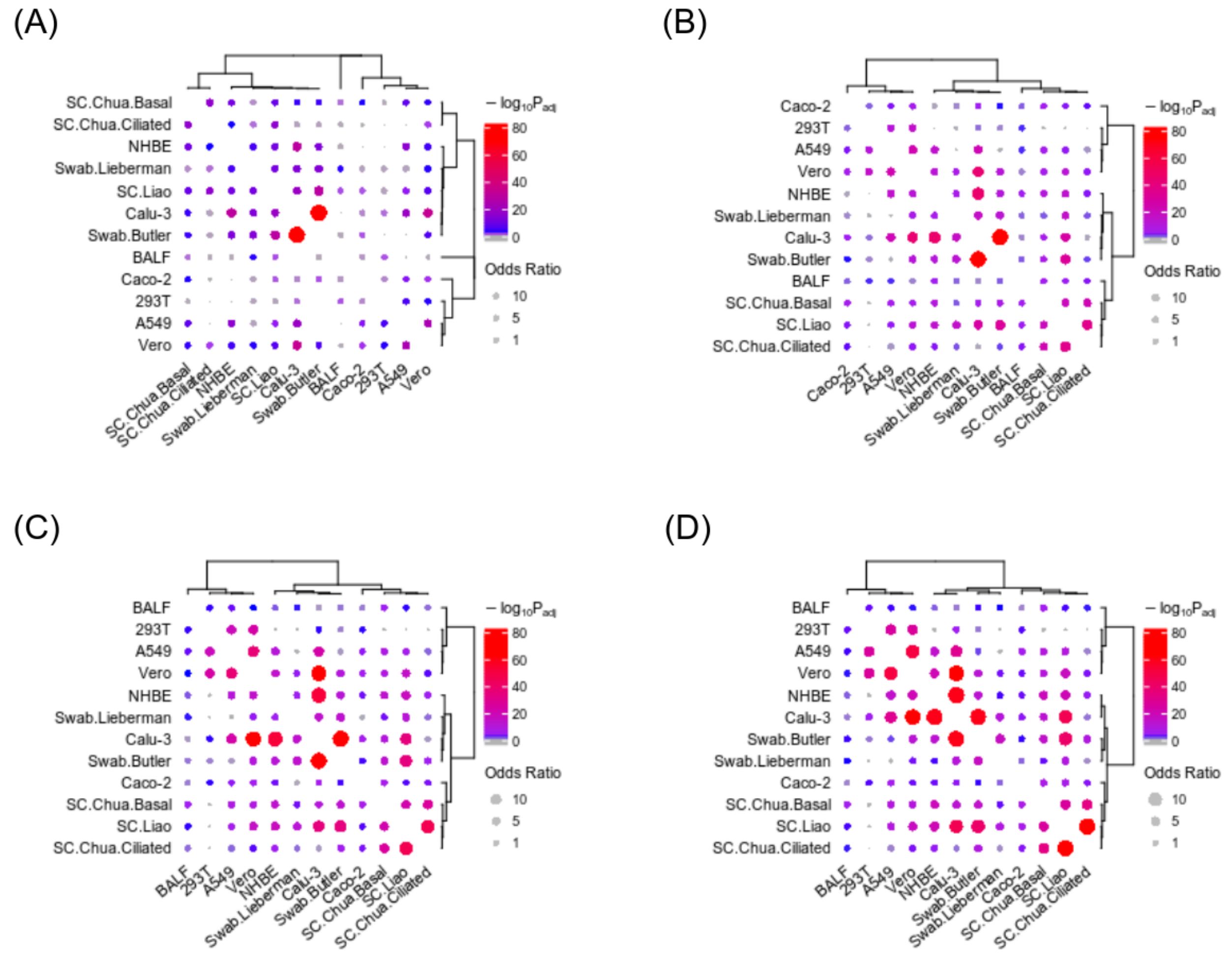


**Appendix Figure S1. Comparison of the differentially expressed (DE) genes upon SARS-CoV-2 infection across datasets with different DE cutoffs.** Visualization of the overlap of the top n=100 **(A)**, 200 **(B)**, 300 **(C)** and 400 **(D)** DE genes between each pair of datasets analyzed using Fisher's exact tests. All the genes up to the top n=400 genes have FDR<0.1 across all datasets. The ranges of odds ratios and P values, as well as the clusterings of the datasets remain similar regardless of the number of top DE genes used. The dot size corresponds to the effect size of the overlap as measured by odds ratio, and the color corresponds to the negative log10 adjusted one-sided P value (grey means below 0.05).

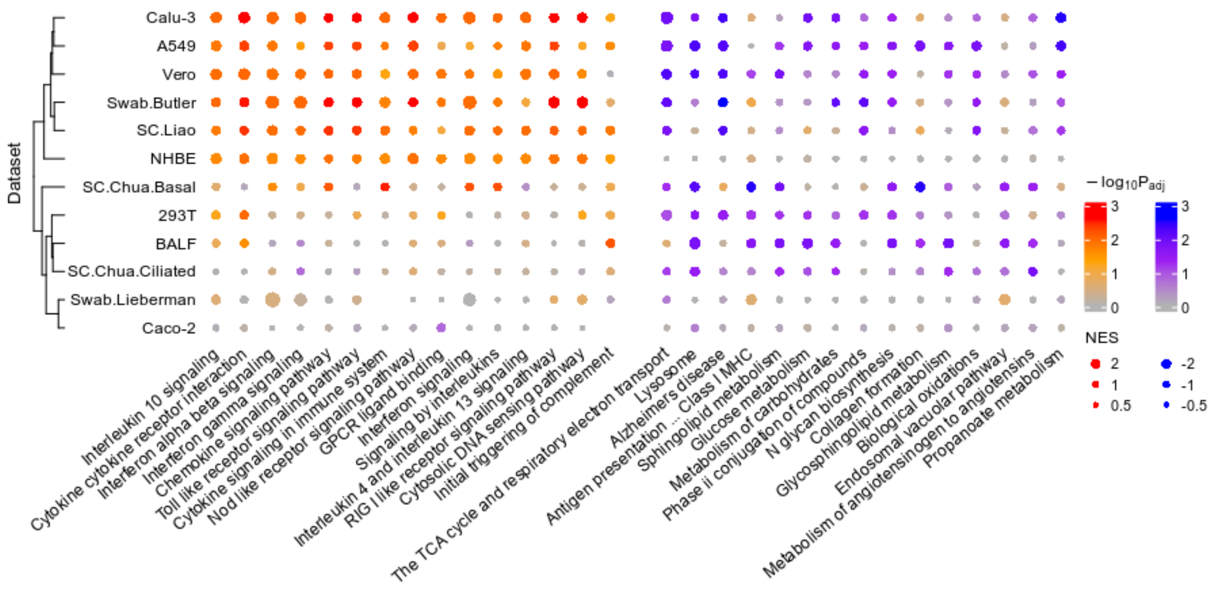


**Appendix Figure S2. Pathway enrichment analysis of SARS-CoV-2-induced differential changes with the alternative differential expression (DE) algorithm limma-voom.** A summary visualization of the GSEA result for the top consistently altered pathways during SARS-CoV-2 infection across the datasets, with more importance given to the various *in vivo* patient datasets (Methods). Unlike Figure 1D in the main text where the bulk RNA-seq dataset DE results were obtained using mostly DESeq2 mixed with other algorithms, here limma-voom was used for the DE analysis of all bulk RNA-seq datasets. The dot color corresponds to the negative log10 adjusted P values from GSEA, with two sets of colors (red-orange and blue-purple) distinguishing up-regulation from down-regulation (positive or negative normalized enrichment scores, i.e. NES); dot size corresponds to the absolute value of NES measuring the strength of enrichment. The left and right-hand side blocks represent the pathways that tend to be consistently up-regulated and down-regulated in infected vs control samples, respectively; within each block, the pathways are ordered by negative sum of log P values across datasets (i.e. Fisher's method).
